# Age- and amyloid-β-dependent initiation of neurofibrillary tau tangles: NLFTau^m/h^, an improved mouse model of Alzheimer’s disease without mutations in *MAPT*

**DOI:** 10.1101/2024.11.04.621900

**Authors:** Sneha Desai, Jiongqiao La, Aya Balbaa, Elena Camporesi, Gunnar Brinkmalm, Kritika Goyal, Argyro Alatza, Jack I. Wood, Takshashila Tripathi, Sumi Bez, Nazar Stasyuk, Haady B. Hajar, Takashi Saito, Takaomi C. Saido, Henrik Zetterberg, John Hardy, Jeffrey N. Savas, Dervis A. Salih, Damian M. Cummings, Jörg Hanrieder, Frances A. Edwards

## Abstract

The lack of models in which ageing and amyloid-β (Aβ) initiate neurofibrillary tau tangles, independent of *MAPT* mutations, limits mechanistic insights into Alzheimer’s disease. Here, we characterise NLFTau^m/h^, a novel mouse model that recapitulates the gradual transition from Aβ accumulation to tau pathology by 24 months. We show that combined effects of ageing, Aβ, and human tau expression promote tau pathology. However, either Aβ plaques or human tau alone can initiate tau pathology, but only in even older mice. This may be exacerbated by increased neuronal vulnerability across all genotypes with age. Moreover, multi-omic analyses reveal that human tau can induce early alterations in mitochondrial and synaptic pathways, priming this latent vulnerability preceding tau pathology. Together, this suggests that ageing and other interacting triggers accelerate the onset of tau pathology. This opens possibilities for translatable testing of improved therapies for Alzheimer’s disease, breaking the link between Aβ and tau pathologies.

## INTRODUCTION

Animal models of Alzheimer’s disease (AD) are essential for understanding disease mechanisms and facilitating drug discovery. Yet, their reliability is often questioned due to limited therapeutic success in humans. Several factors may contribute to these translational gaps.

Most notably, AD is only clinically diagnosed due to the cognitive decline that follows the development of neurofibrillary tau tangles (NFTs), which occurs decades after the initial accumulation of amyloid-β (Aβ) plaques in the human brain. Most commonly, diagnosis occurs in late old age, approximately 81- 86 years^1^. Moreover, more than 95% of AD cases are sporadic^2^, whereas most animal models rely on familial mutations, possibly limiting their relevance to the broader patient population. Therefore, to understand the initiation and progression of Aβ and NFT pathology, an ideal model would spontaneously develop early Aβ pathology and later, in old age, NFTs and neurodegeneration, along with associated cognitive deficits. The only such models reported to develop AD-like pathology naturally, the Chilean degus^3^ and the marmoset^4^, are valuable but they require expensive colonies and only a subset of animals develop pathology with many years of ageing, limiting their practicality.

Existing mouse models reproduce the pre-clinical stages of plaque pathology; however, they often also rely on transgenic overexpression technology and despite overexpression and rapid onset of heavy plaque loads, they do not go on to develop NFTs. In fact, NFTs have only previously been initiated in mice that bypass any link between Aβ and tau pathology by the inclusion of mutations in *MAPT,* the gene encoding the tau protein^5,6^. Endogenous murine tau seems to be resistant to misfolding or aggregation under physiological conditions, unless seeded by pathological tau, suggesting a species-specific difference in templating^7^. While the *MAPT* sequence in adult mice and humans is similar, there are important differences in the isoforms expressed, likely affecting folding^8^, and in the N-terminal region, affecting a range of binding partners^9,10^.

To address these limitations, recent efforts have focused on humanising tau to capture human tau biology more closely. Humanised *MAPT* knock-in mice recapitulate human-specific splice isoform ratios and show improved modelling of tau propagation compared to wildtype (WT) counterparts^11^. Importantly, these models avoid artifacts associated with transgene overexpression and genome disruption^12–14^.

Substantial progress has been made in understanding tau biology, with studies demonstrating that NFTs are associated with the mislocalisation of soluble tau from axons to somatodendritic compartments, where it accumulates in a hyperphosphorylated and aggregated form that is strongly associated with neurodegeneration^14–16^. Moreover, a recent study has reported that different *MAPT* mutations exhibit heterogeneity in the trajectories of tau pathogenesis, illustrating how distinct triggers can drive divergent mechanisms^17^. However, despite these advances, the upstream mechanisms by which Aβ pathology is suggested to drive NFT formation remain poorly understood, hindering efforts to develop therapies that prevent progression to clinical stages. Hence, there is an urgent need to improve existing mouse models by addressing several key factors:

1. Rising Aβ deposition should induce the formation of NFTs, independent of mutations not relevant to AD.
2. A delay should occur between the onset of Aβ plaque accumulation and the establishment of tau pathology.

In this study, we introduce a model in which an amyloid mouse is crossed with a heterozygous humanised *Mapt* mouse. Both these knock-in models were previously developed in RIKEN, Japan^11,18^. In this novel mouse, NLFTau^m/h^, the interaction of ageing, Aβ pathology, and human tau can initiate NFTs without the introduction of *MAPT* mutations. Interestingly, we also find that plaques or human tau alone can initiate NFTs in mice. However, in the absence of their combination, NFTs are only initiated at extreme old age (28 months), when mortality approaches 50%, limiting experimental utility. Combining the slow development of plaques into old age in *App*^NL-F/NL-F^ mice with the presence of human tau accelerates the initiation of NFTs to 24 months of age, when most mice can still be studied. Our multi-omics analyses reveal that human tau expression induces early, network-level alterations in mitochondrial and synaptic pathways, suggesting a primed stage of vulnerability preceding tau pathology. Collectively, our findings suggest that ageing establishes a baseline vulnerability, while the interaction of Aβ plaques and human tau lowers the threshold for the initiation of NFTs. The NLFTau^m/h^ model therefore provides a physiologically relevant platform to probe the mechanistic underpinnings of the interaction of Aβ-dependent tau pathology in AD. This is essential for development of therapeutic strategies aimed at preventing the clinical manifestation of tau pathology.

## RESULTS

Resistance of mice to forming NFTs could arise from differences between human and mouse in the tau N-terminal region and isoform expression (Fig. S1) ^18^. Hence, to align the slow time-course of plaque development in sporadic AD and include the important factor of ageing, we have developed a model, starting with the *App^NL-F/NL-F^* (NLF) knock-in mouse^19^ from 24 months of age. Like in sporadic AD in humans, NLF mice show a gradual plaque onset from mid-life, progressively increasing through old age^19,20^. The NLF mouse was crossed with a knock-in humanised *Mapt* mouse^11^, such that the *App* mutations were homozygous and the humanised *Mapt* heterozygous (NLFTau^m/h^). This introduces humanised tau, while minimising potential developmental artefacts that could arise from completely removing murine tau.

### Comparable tau protein profiling patterns are observed in NLFTau^m/h^ and human AD

Initially, we measured the relative abundance of different tau isoforms in cortical tissue. Using both western blot and immunoprecipitation, combined with liquid-chromatography mass spectrometry (IP-MS), the relative abundance of tau peptides in 24-month-old WT, NLF, Tau^m/h^ and NLFTau^m/h^ mice were compared. As expected, adult WT and NLF mice lacked the 3 carboxyl-terminal repeat domain (3R) isoforms of tau^21^, whereas Tau^m/h^ and NLFTau^m/h^ exhibited all six isoforms (Fig. 1A, C). Both methods confirmed that the 3R:4R isoform ratio in both Tau^m/h^ and NLFTau^m/h^ mice resembled the approximately equal levels reported in humans, both in healthy controls and AD patients^22^(Fig. 1B, D).

**Figure 1.**
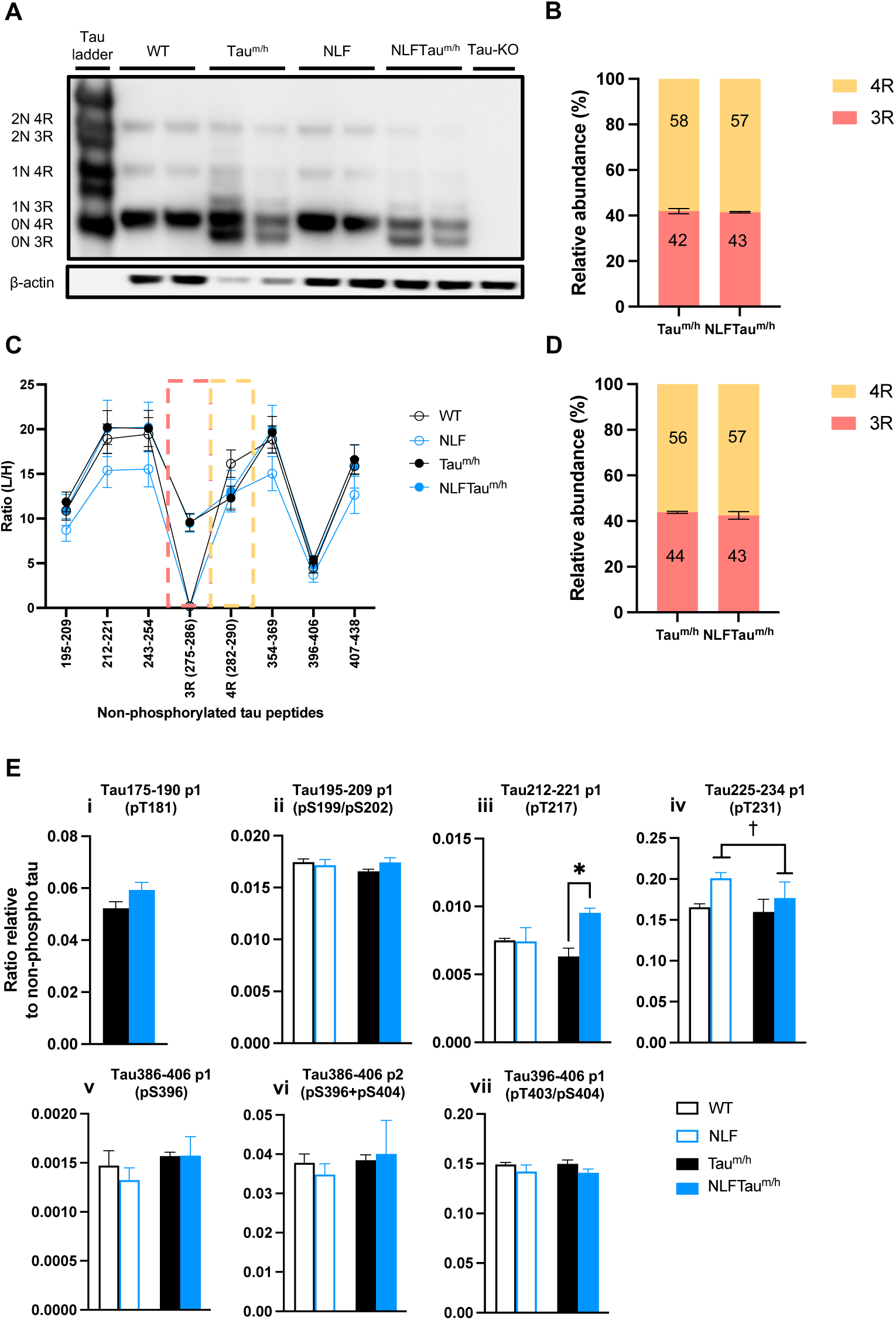
Analysis of tau peptides in 24-month-old WT, NLF, Tau^m/h^ and NLFTau^m/h^ mice. **(A)** Immunoblotting of total tau detected by Tau-5. β-actin is used as a loading control. **(B)** Relative quantification of 3R and 4R tau isoforms from **(A)** (Tau^m/h^ n = 4, NLFTau^m/h^ n = 3 mice). **(C)** Quantification of non-phosphorylated tau peptides by IP-MS analysis (Two-way ANOVA; WT n = 4, NLF n = 5, Tau^m/h^ n = 4, NLFTau^m/h^ n = 3 mice). **(D)** Relative abundance of 3R and 4R tau isoforms from **(C)**. **(E)** Abundance of tau peptides carrying one (p1) or two (p2) phosphorylations, relative to non-phosphorylated 354-369 tau peptide level by IP-MS analysis (Two-way ANOVA; WT n = 4, NLF n = 5, Tau^m/h^ n = 4, NLFTau^m/h^ n = 3 mice). *Post hoc* Sidak’s multiple comparisons test shown as *p<0.05, main effect of *App* mutations shown as †p<0.05. Data in **B-E** is represented as mean ± SEM. (**E i)** As the standards used were for human tau, for phosphorylation sites within sequence sections that differ between mouse and human only genotypes carrying human tau were analysed.

We next examined different non-phosphorylated and phosphorylated tau residues. While non-phosphorylated tau peptides, in regions analogous between mouse and human tau, were unchanged across genotypes (Fig. 1C, Table S1), several changes were evident in phosphorylated tau residues (Fig. 1E, Table S1). The largest difference in phosphorylation detected was in the singly phosphorylated (p1) tau peptide 212-221, being significantly more abundant in NLFTau^m/h^ mice than age-matched controls (Fig. 1Eiii). Tandem mass spectrometry analysis revealed that the primary phosphorylation site was pT217. The pT231 residue was also significantly higher in both NLF and NLFTau^m/h^ mice compared to the other genotypes (Fig. 1Eiv). Both these residues are particularly notable, being relevant to human fluid biomarkers of AD^23–25^.

### Humanisation of tau accelerates the onset of tau hyperphosphorylation in pyramidal cell somas

We went on to investigate p-tau pathology in the soma of hippocampal pyramidal neurones, which would be indicative of initial NFT formation^23,26^. Specifically, we used immunohistochemistry to quantify tau phosphorylation at the serine396 epitope (pS396), a site commonly phosphorylated within the paired helical filaments of NFTs^27^. Although changes in this or other C-terminal p-tau sites were not detected in bulk tissue analysis (Fig. 1Ev-vii), it is likely, especially at early stages, that spatial analysis would be necessary to detect the initiation of NFTs.

As tau protein is normally primarily located in axons, the onset of tau pathology in the soma of pyramidal neurones would be expected to increase relative to axons. To assess this, we measured pS396 florescence intensity in the pyramidal cell body layers of CA1 and CA3 hippocampal regions and compared it with intensity in adjacent axon-rich regions (Fig. 2A). Analysis of the intensity ratio of pS396 staining in soma to axon regions revealed an increase in pS396 protein ratio in both CA1 and CA3 hippocampus in 24-month-old NLFTau^m/h^ mice compared to age-matched controls (Fig 2B, D). At this age, a small proportion of neurones in some WT, Tau^m/h^ and NLF mice also showed a shift of pS396 staining to the soma, although this was greatest and more consistent in NLFTau^m/h^ mice in both regions. Heatmaps were generated based on the intensity ratio in CA1 and CA3 hippocampal regions (Fig. 2C, E). Note that a ratio of 1:1 means that the intensity of pS396 staining was equal in the somatic and axonal regions while higher ratios indicate a distribution more towards the soma.

**Figure 2.**
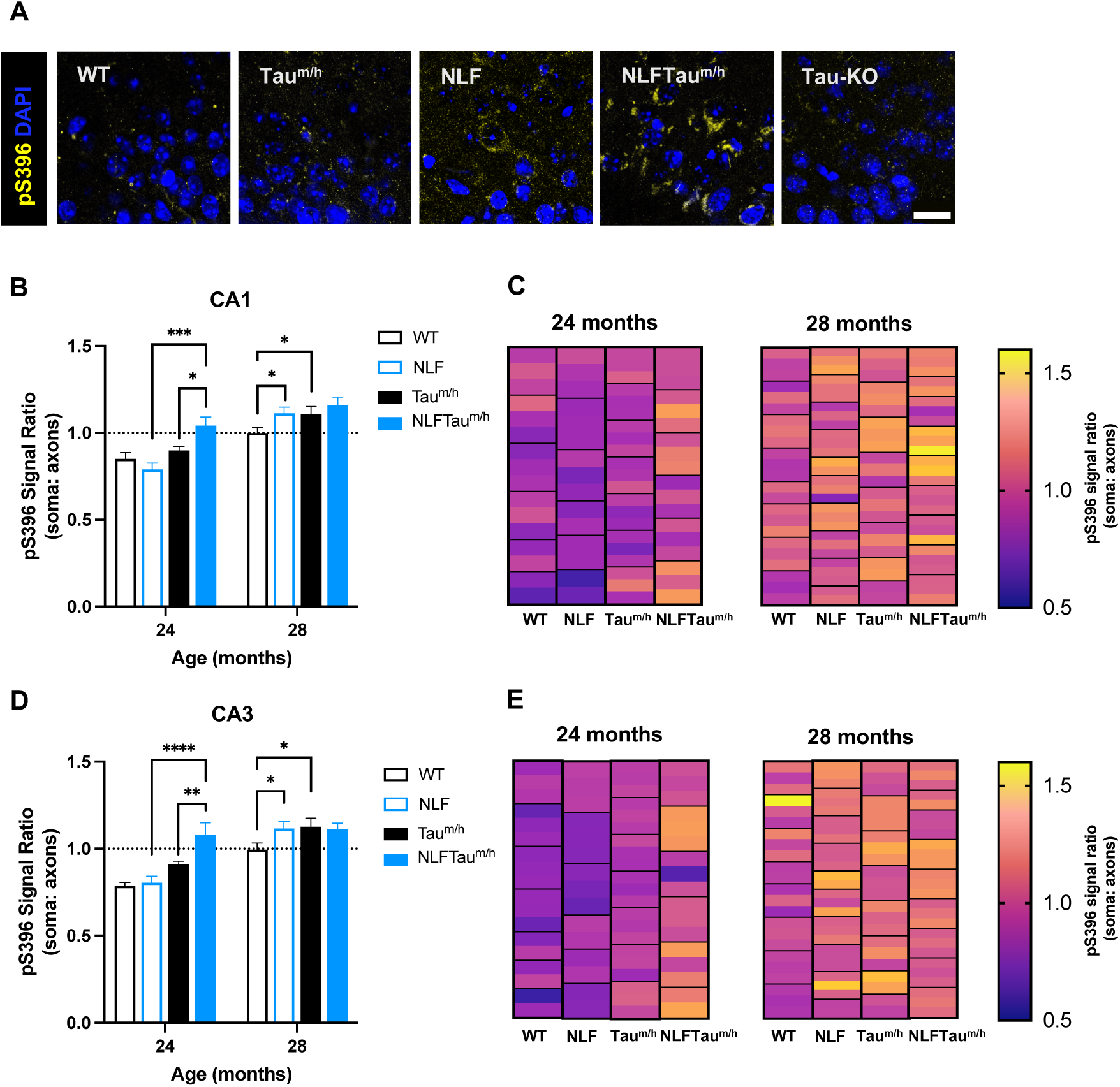
Changes in distribution of pS396 staining across age and genotype in hippocampal neurones. **(A)** Representative images of pS396 staining across genotypes in CA1 hippocampal neurones at 24 months. Scale bar: 20 μm. **(B, D)** pS396 florescence intensity in CA1 and CA3 hippocampal pyramidal neurones as a ratio of intensity in the neuronal soma versus the axonal region across age and genotype. Generalised linear mixed model analysis followed by a least significant difference *post hoc* analysis: ****p<0.0001, ***p<0.001, **p<0.01, *p<0.05. Note that *post hoc* tests only assess differences within one age between genotypes with one different variable. Across age *post hoc* analyses: For **(B)** WT: p<0.05, NLF: p<0.0001, Tau^m/h^: p<0.001, NLFTau^m/h^: p <0.05. **(D)** WT: p<0.01, NLF: p<0.0001, Tau^m/h^: p<0.001. Data in B and D are presented as mean ± SEM. **(C, E)** Heatmaps show the distribution of pS396 florescence intensity across age and genotype in CA1 and CA3 hippocampal pyramidal neurones. Each black bold box represents a separate mouse, while the boxes within indicate technical replicates: WT n = 6-8, NLF n = 6-10, Tau^m/h^ n = 7-8, NLFTau^m/h^ n = 6-9 mice (2-3 technical replicates per mouse).

While the intensity ratio of soma to axonal pS396 clearly indicated a more advanced initiation of tau pathology in the NLFTau^m/h^ mice at 24 months, compared to other genotypes, the ratio was variable between mice, and so we investigated whether evidence of initiation of tau tangles would be more consistent in even older mice. Interestingly, by 28 months, although the ratio increased and was more consistent in the NLFTau^m/h^ mice, compared to the ratios at 24 months, all genotypes showed an age-related increase in pS396 ratios. In both hippocampal regions, pS396 ratio in 28-month-old NLF and Tau^m/h^ mice increased to match the relative intensity observed in NLFTau^m/h^ mice at this age. Although WT mice also showed an age-related increase, the overall ratio (∼1) remained lower compared to other genotypes. Together, these findings suggest that while human tau, in the presence of Aβ plaques, accelerates the onset of tau hyperphosphorylation in the soma of hippocampal pyramidal cells, ageing independently promotes AD-related p-tau pathology, even when either human tau or Aβ pathology is present alone.

### Humanisation of tau also promotes accumulation of hyperphosphorylated tau within cored plaques

Amyloid mouse models generally show p-tau within and in the immediate vicinity of plaques, putatively in dystrophic neurites^19,28^. Thus, having established the acceleration of the initiation of NFT initiation in NLFTau^m/h^ mice, dependent on age and the presence of Aβ plaques, we went on to investigate whether the inclusion of human tau would also affect the deposition of plaques and p-tau levels in the immediate vicinity of these plaques.

Plaques can be classified according to their Aβ peptide signatures and morphology, using a combination of two luminescent conjugated oligothiophene dyes (LCOs), q- FTAA and h-FTAA (Fig. S2A), together with an Aβ antibody, 6E10. q-FTAA preferentially binds to mature β-pleated amyloid fibrils, whereas h-FTAA binds to all β-pleated amyloid fibrils^29,30^. In contrast, 6E10 binds to all forms of Aβ^31^. We could thus distinguish three categories (Fig. 3A):

1. q-FTAA+ve, h-FTAA+ve, Aβ+ve (cored plaques)
2. q-FTAA-ve, h-FTAA+ve, Aβ+ve (fibrillar plaques)
3. q-FTAA-ve, h-FTAA-ve, Aβ+ve (diffuse Aβ).

**Figure 3.**
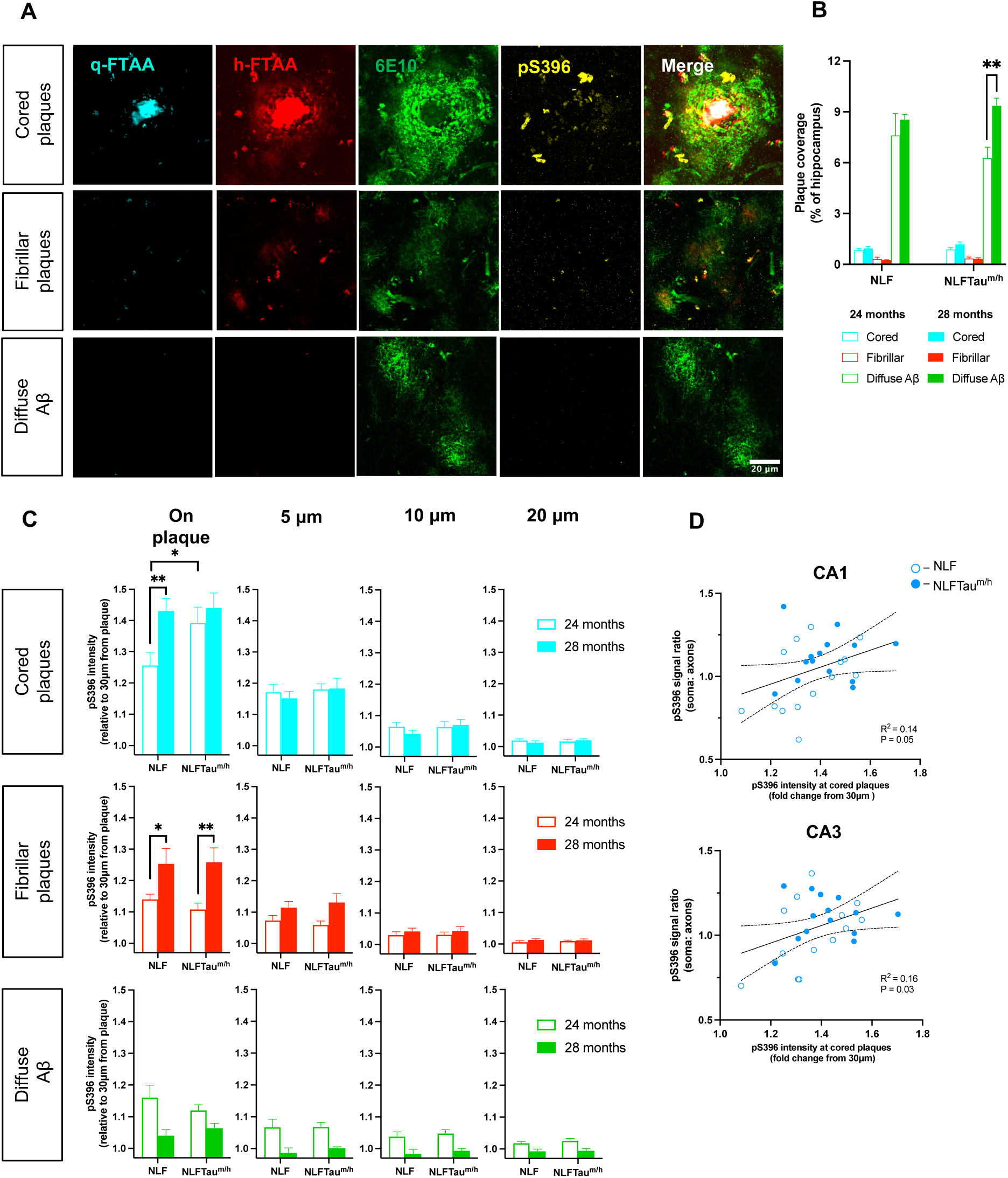
Distribution of pS396 staining surrounding different plaque types across age and genotype. **(A)** Representative images of different plaque types and associated pS396 protein levels. **(B)** Hippocampal coverage of different plaque types in 24- and 28-month-old NLF and NLFTau^m/h^ mice. Generalised linear mixed model analysis followed by a least significant difference *post hoc* analysis. **p<0.01. **(C)** Mean pS396 tau fluorescence pixel intensity on the plaque expressed as a fold change relative to 30 μm away from the plaque in 24- and 28-month-old NLF and NLFTau^m/h^ mice. Linear mixed-effects model with genotype, age, distance and plaque type as fixed effects, accounting for repeated measures within animals. Genotype x age x plaque type interactions were first assessed at specific distances, followed by analysis of the genotype x age interaction within each plaque type at the “on plaque” distance only, as other distances showed no significant effects. *Post hoc* comparisons were performed using least significant difference: *p<0.05; **p<0.01. Data in **B,C** are presented as mean ± SEM. NLF N = 6-10 mice; NLFTau^m/h^ N = 6-9 mice (2-3 technical replicates per mouse). **(D)** Correlation of pS396 intensity within CA1 and CA3 neuronal soma versus pS396 intensity at cored plaques in 24- and 28-month-old NLF and NLFTau^m/h^ mice. Lines represent linear regression with 95% confidence intervals.

In both 24- and 28-month-old mice, a significantly greater hippocampal area was occupied by diffuse Aβ compared to cored and fibrillar plaques (Fig. S2B). While the diffuse plaque coverage was consistent in the NLFTau^m/h^ mice, increasing significantly with age, the coverage in NLF mice was more variable but not significantly different from the NLFTau^m/h^ mice (Fig. 3B).

p-Tau (pS396 levels) staining was punctate, increasing with proximity to Aβ in all cases, with the greatest increase in the immediate vicinity of cored plaques. At 24 months old, this increase in the vicinity of cored plaques was greater in NLFTau^m/h^ mice than in NLF mice. However, by 28 months, both genotypes increased pS396 signal compared with 24 months and displayed a similar distribution of signal intensity. Thus, as observed in the soma, the maximum pS396 response measured was similar between genotypes by 28 months, although this response occurred earlier in NLFTau^m/h^ mice. In fibrillar plaques, the pS396 signal was lower than in cored plaques, however the response was similar between genotypes at both ages, with both increasing from 24 to 28 months (Fig. 3C). Interestingly, little, if any, pS396 signal was observed surrounding diffuse Aβ, and this pattern remained consistent across genotypes and ages, suggesting a differential effect of different plaque types on p-tau pathology.

Considering the similar pattern of development of p-tau accumulation in cored plaques and in the soma at these ages, we investigated whether the changes in these two locations correlated across individual mice. We observed a positive correlation between the strength of pS396 staining in cored plaques and in the hippocampal soma of individual NLF and NLFTau^m/h^ mice, in both CA1 and CA3 regions (Fig. 3D). Collectively, these results align with our earlier findings, indicating that human tau accelerates tau hyperphosphorylation within cored plaques, a process that is also independently enhanced by ageing and may be linked to NFT initiation.

### NLF and NLFTau^m/h^ mice show reduced survival without significant neurodegeneration

Neurodegeneration and reduced life span are pathological hallmarks of late-stage AD^32,33^. To assess whether survival differed between WT, NLF, Tau^m/h^ and NLFTau^m/h^ mice, we generated Kaplan-Meier survival curves. Using a Cox proportional hazards model, we found that survival probability was significantly reduced in mice carrying *App* mutations, but with no effect of human tau (Fig. 4A). Consistent with this, other amyloid mouse models also exhibit reduced survival compared to WT mice^34^. However, this reduction in survival was not reflected in the number of hippocampal neuronal pyramidal cells in both CA1 and CA3 regions (Fig. 4B-C). Note that while around 80% of mice survived to 24 months, the number of animals surviving dropped rapidly with ≤ 50% by 28 months. This suggests that while these models depict NFT initiation, they do not reach end-stage AD within their lifetime.

**Figure 4.**
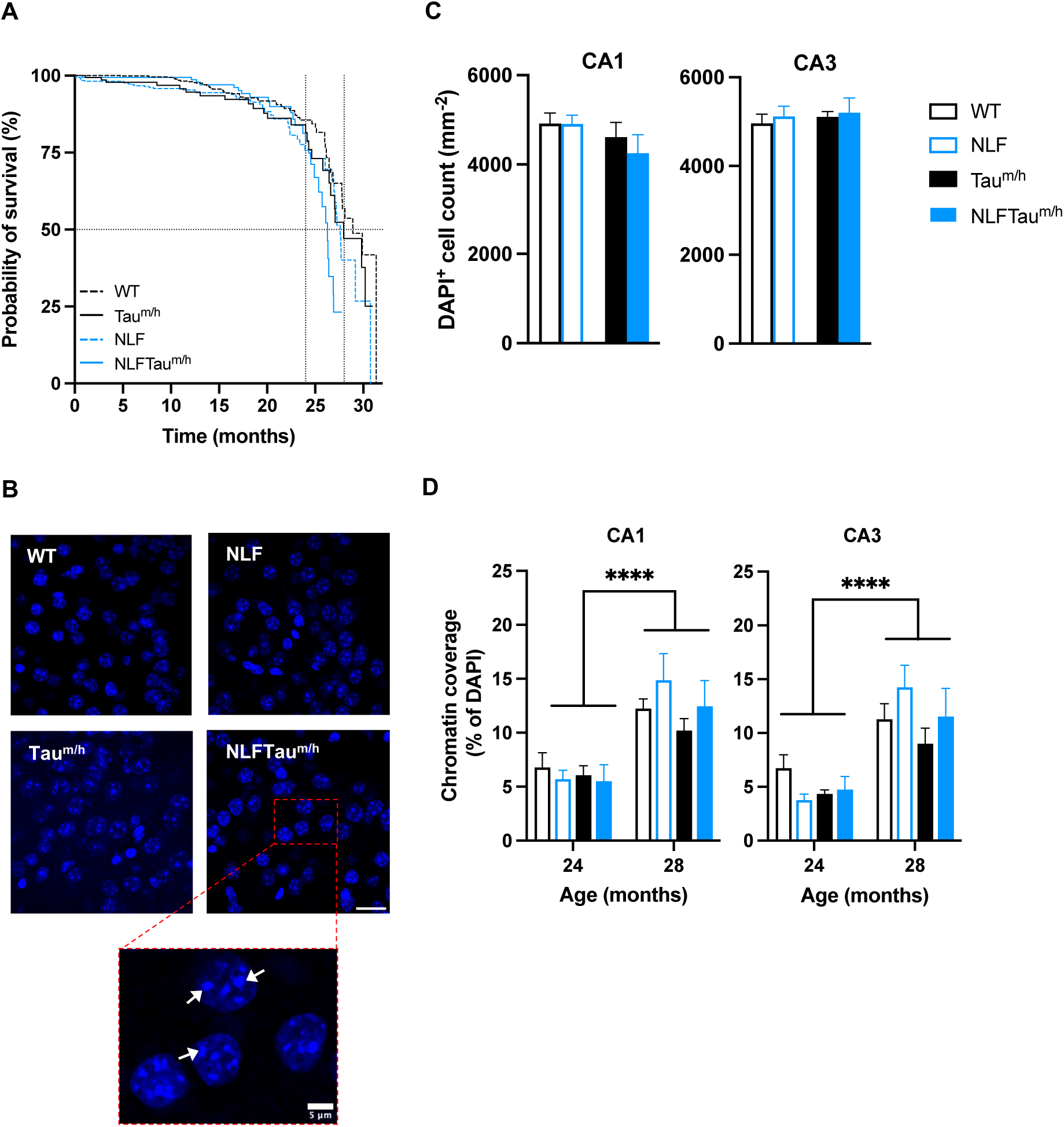
Age- and genotype-dependent decline in lifespan and neuronal health. **(A)** Kaplan-Meier survival curves with statistical significance determined using a Cox proportional hazards model showing a main effect of *App*: *\*\**p<0.01 (WT = 864, Tau^m/h^ = 172, NLF = 510, NLFTau^m/h^ = 186 mice). **(B)** Representative images of DAPI-stained nuclei of CA1 pyramidal neurones in 24-month-old mice. Scale bar: 20 μm. In-set shows condensed chromatin within nuclei; examples indicated by arrows. **(C)** DAPI-stained nuclei counts in CA1 and CA3 pyramidal neurones at 28 months across WT, NLF, Tau^m/h^ and NLFTau^m/h^ mice. **(D)** Chromatin coverage in DAPI-stained nuclei of hippocampal neurones within CA1 and CA3 regions at 24 and 28 months. Generalised linear mixed model analysis showing a main effect of age ****p<0.0001. Data in **C, D** are presented as mean ± SEM. WT = 5-8, Tau^m/h^ = 6-8, NLF = 6-8, NLFTau^m/h^ = 5 mice (2-3 technical replicates per mouse).

Chromatin condensation within neuronal nuclei has been reported to be an early indicator of apoptosis^35,36^ and can be quantified as a measure of apoptotic progression. This process could influence differences in tau pathology progression or neuronal survival. Therefore, we measured the average coverage of compacted chromatin per nucleus in CA1 and CA3 regions (Fig. 4B). Compacted chromatin coverage increased from 24 to 28 months but was not significantly different between genotypes (Fig. 4D). These findings suggest that the increase in chromatin condensation reflects an age-related vulnerability of neurones rather than an effect driven by Aβ or tau pathology.

### Proteomic analyses identify mitochondrial and synaptic alterations in NLFTau^m/h^ mice

To gain insights into molecular mechanisms underlying tau hyperphosphorylation and its interaction with Aβ pathology with or without human tau, we performed proteomic analyses in whole cortical tissue of WT, Tau^m/h^, NLF, and NLFTau^m/h^ mice. Analysis was carried out in 24-month-old mice when somatic p-tau levels differed between genotypes.

We used liquid chromatography-mass spectrometry proteomic profiling, identifying and quantifying 3624 proteins across all animals. We analysed the protein levels using limma and generated volcano plots to visualise genotype-specific comparisons (Fig. 5Ai-iv). NLF versus WT and NLFTau^m/h^ versus Tau^m/h^ comparisons, which capture Aβ-associated changes, yielded only a few differentially expressed proteins (DEPs). Notably, some of these proteins, including apoE and COL25A1, are encoded by allelic variants that have been associated with increased risk of AD^37,38^. In contrast, Tau^m/h^ versus WT and NLFTau^m/h^ versus NLF, which assess human tau-associated effects, yielded 32 and 92 DEPs respectively. These results indicate more extensive proteomic alterations in NLFTau^m/h^ than in NLF or Tau^m/h^ mice versus their controls, highlighting that the expression of human tau resulted in more significant changes when compared to endogenous mouse tau. Complete comparisons can be found in Supplementary Table S2.

**Figure 5.**
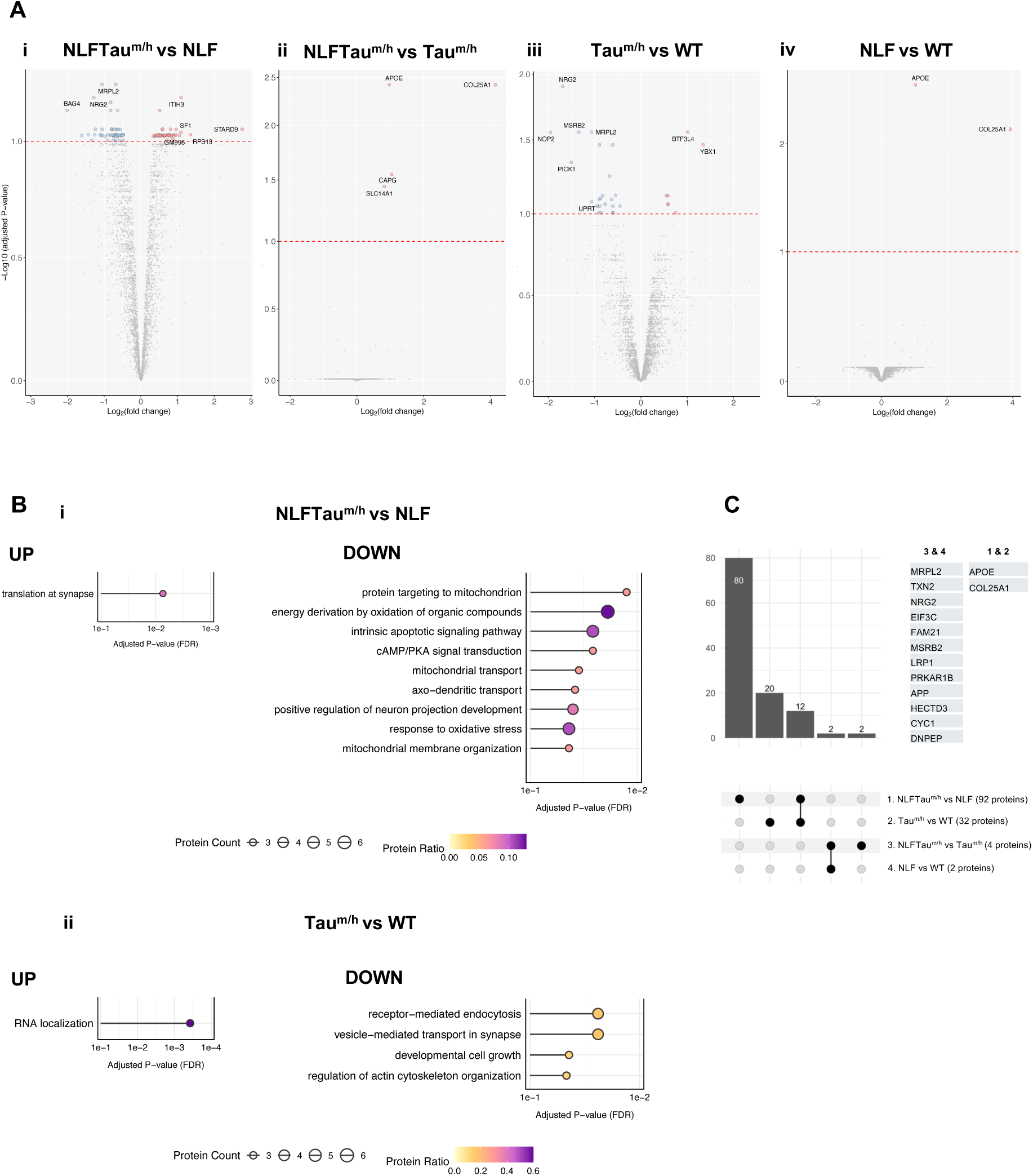
Bulk proteomics analysis in 24-month-old WT, NLF, Tau^m/h^ and NLFTau^m/h^ cortex. **(A)** Volcano plots displaying DEPs in **(i)** NLFTau^m/h^ vs NLF, **(ii)** NLFTau^m/h^ vs Tau^m/^h, **(iii)** Taum^/h^ vs WT, and **(iv)** NLF vs WT. The dashed red line indicates an adjusted p-value of 0.1, above which proteins are considered significant. Labelled proteins are those with absolute log_2_(fold-change) > 1. **(B)** Overrepresentation GO analysis of up- and down-regulated DEPs (adj. p-values) from NLFTau^m/h^ vs NLF in **(i)** and Tau^m/h^ vs WT in **(ii)**, highlighting top biological processes (BP). **(C)** UpSet plot showing shared DEPs across comparisons seen in **(A)**, with shared proteins listed. WT = 4, Tau^m/h^ = 4, NLF = 4, NLFTau^m/h^ = 4 mice.

We performed gene ontology (GO) analysis on up- or downregulated DEPs from NLFTau^m/h^ versus NLF (Fig. 5Bi) and Tau^m/h^ versus WT (Fig. 5Bii). In NLFTau^m/h^ mice, downregulated proteins were enriched in processes related to mitochondrial function and transport, including proteins such as BID, CPT2, MSRB2, and TRAP1. In contrast, upregulated proteins were enriched in synaptic translation, driven by ribosomal proteins such as RPL4, RPL7A, and RPS13. In Tau^m/h^ mice, downregulated proteins were enriched in processes linked to regulation of synapse structure and actin cytoskeletal organisation. Upregulated proteins, specifically, HNRNPAB, SARNP, and YBX1, were associated with RNA localisation. All enriched terms and associated proteins can be found in Supplementary Table S3. Notably, some DEPs overlapped across multiple comparisons (Fig. 5C), suggesting that certain proteins are commonly differentially expressed due to either humanised tau or Aβ pathology. Additional proteins were dysregulated only under the combined presence of Aβ burden and human tau. One such example is the microglial protein CAPG, which has been consistently reported as increased within both Aβ plaques and NFTs in human AD proteomic studies^39^.

To uncover coordinated patterns of proteome-wide changes, we conducted weighted co-expression network analysis. Hierarchical clustering of protein levels identified ten distinct modules, of which four exhibited genotype-dependent shifts in eigenprotein values in mice expressing human tau (Fig. 6A). While all four modules are shown in Fig. 6A, the red and brown modules displayed the most pronounced changes in NLFTau^m/h^ and Tau^m/h^ relative to NLF and WT mice and were therefore selected for further functional characterisation.

**Figure 6.**
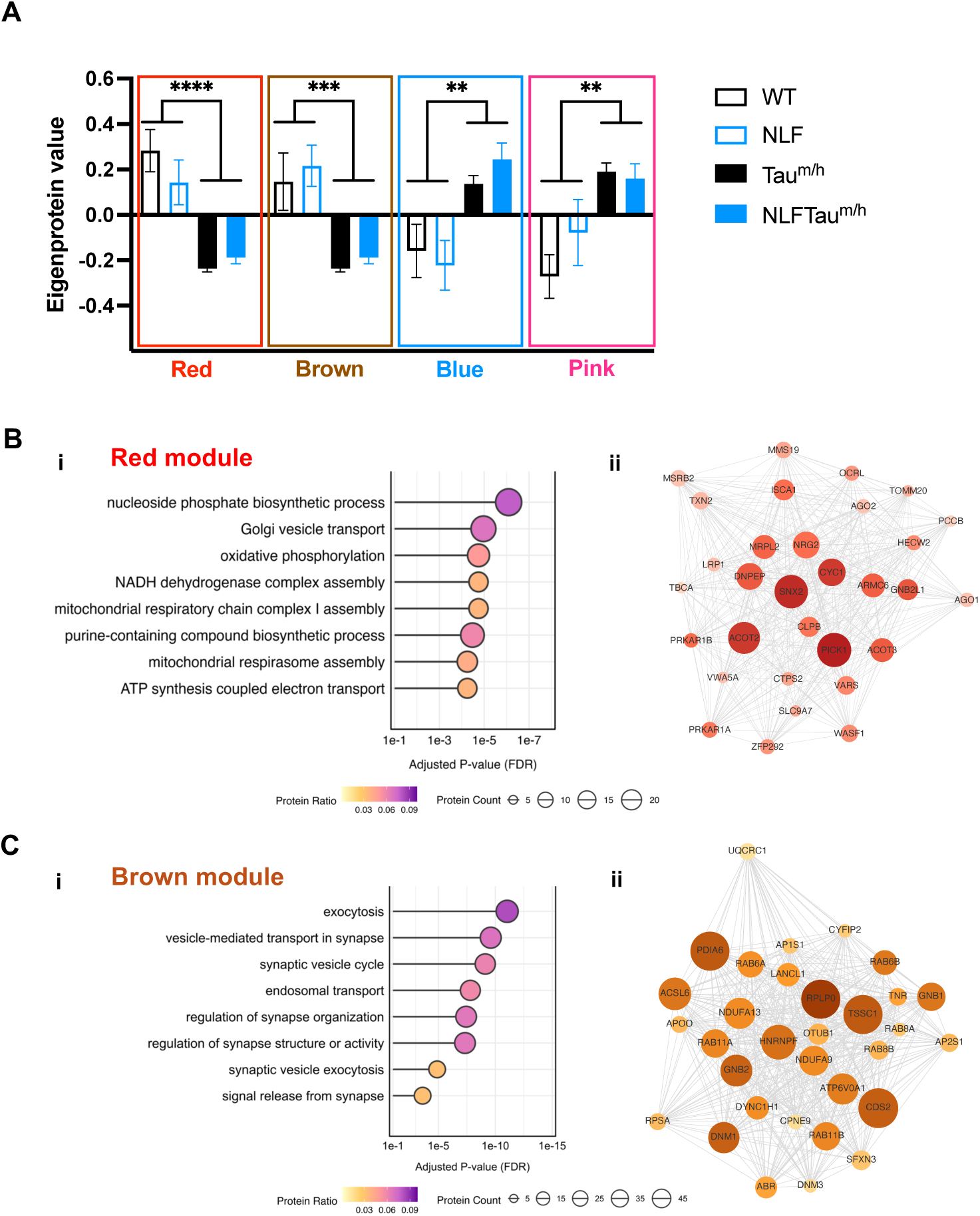
Weighted co-expression network analysis-derived protein modules in 24-month-old WT, NLF, Tau^m/h^ and NLFTau^m/h^ mice. **(A)** Module eigenprotein values separated by genotype. Colours correspond to the assigned module ME colours. Two-way ANOVA showing a main effect of Tau^m/h^: ****p<0.0001, ***p<0.001, **p<0.01. **(B-C) (i)** GO enrichment analysis, highlighting top BP terms identified by overrepresentation analysis with adj. p-value < 0.05. **(B-C) (ii)** Network plot of top 30 proteins. Node size represents connectivity (number of protein-protein interactions). Increasing colour intensity of node indicates higher module membership (kME). WT = 4, Tau^m/h^ = 4, NLF = 4, NLFTau^m/h^ = 4 mice.

GO analysis revealed that the red module was enriched for mitochondrial energy metabolism, oxidative phosphorylation, and oxidative stress response pathways, driven by proteins such as CYC1, ACOT2, and DNPEP (Fig. 6B). The brown module was associated with synaptic vesicle trafficking and structural organisation, featuring proteins including DNM3, GNB2, CYFIP2, and multiple RAB family members (Fig. 6C). These functional profiles are consistent with the enrichment patterns observed in Fig. 5B. A complete list of GO terms enriched within each module and their associated proteins can be found in Supplementary Table S4.

### Targeted transcriptomic analysis reveals mitochondrial and synaptic transcriptional dysregulation in NLFTau^m/h^ mice

Building on the reduction in mitochondrial and synaptic proteins, we next performed targeted transcriptomic profiling to assess whether protein alterations in these functional areas were accompanied by transcriptional dysregulation.

Gene expression counts from hippocampal mRNA were initially corrected and normalised. A subset of 1,063 genes encoding mouse mitochondrial proteins was curated from the MitoCarta 3.0 dataset^40^, and normalised expression values were analysed using the limma-voom pipeline in 24-month-old WT, NLF, Tau^m/h^, and NLFTau^m/h^ mice.

Consistent with the proteomic findings, differential expression was minimal in NLF versus WT or NLFTau^m/h^ versus Tau^m/h^ comparisons. Only *Acadm* and *Cryz* were significantly downregulated in the Tau^m/h^ versus WT comparison. In contrast, the NLFTau^m/h^ versus NLF comparison identified 78 differentially expressed genes (DEGs; Fig. 7A). These genes were predominantly associated with metabolic processes as well as mitochondrial DNA maintenance, translation and oxidative phosphorylation (Fig. 7B), and were largely localised to the mitochondrial matrix. Analysis of the top DEGs across genotypes indicated that many transcriptional changes were associated with the presence of human tau, and the addition of *App* mutations further reduced expression of these genes in NLFTau^m/h^ mice (Fig. 7C). Complete comparisons and their associated functions and sub-mitochondrial localisation can be found in Supplementary Table S5 and S6.

**Figure 7.**
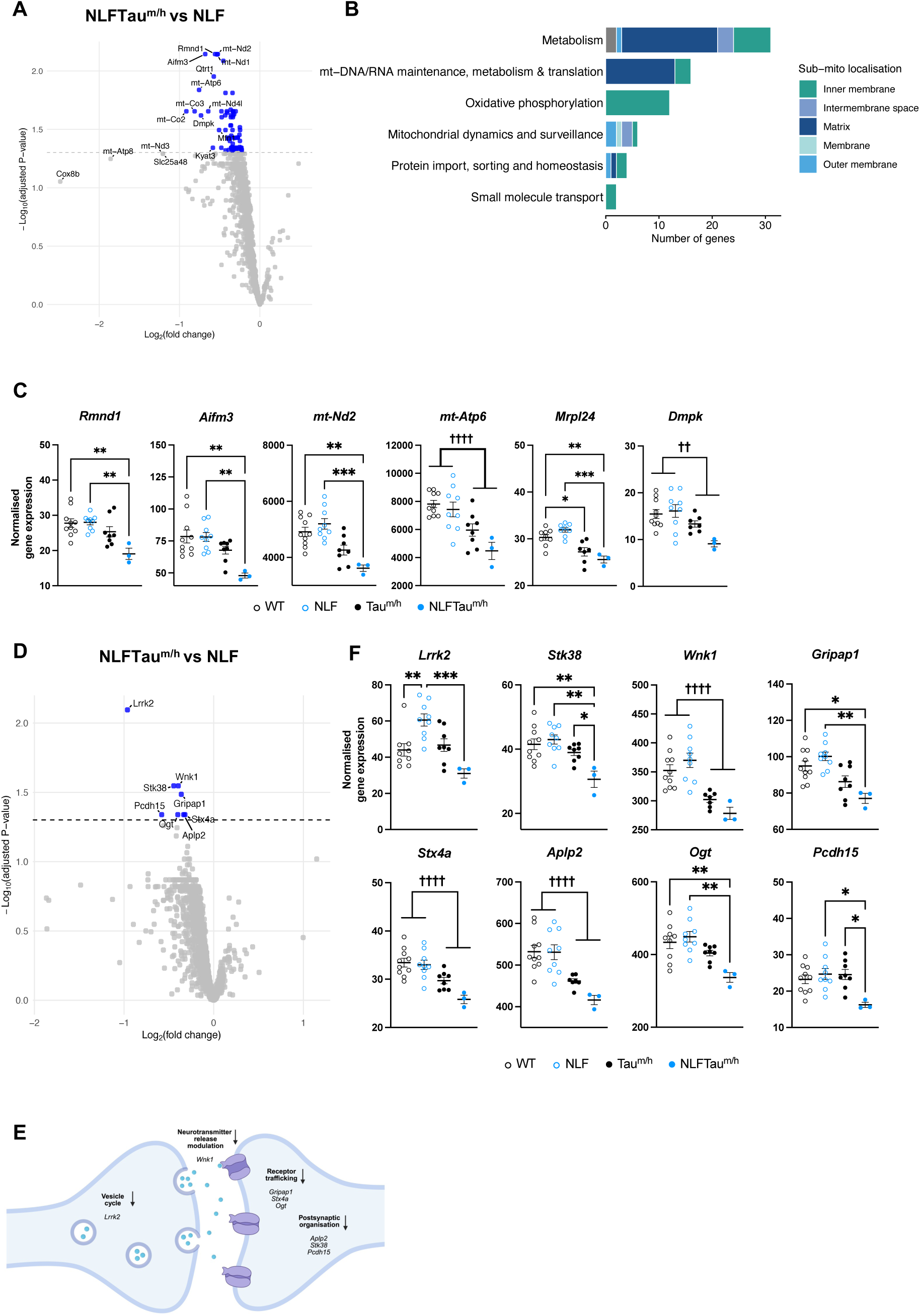
Mitochondrial and synaptic gene expression profiling in 24-month-old WT, NLF, Tau^m/h^ and NLFTau^m/h^ mice. **(A)** Volcano plot displaying DEGs in NLFTau^m/h^ vs NLF, derived from the MitoCarta 3.0 mitochondrial gene set. The dashed grey line indicates an adjusted p-value threshold of 0.05, above which genes are considered significant. Labelled genes have an absolute log₂(fold change) greater than 0.5. **(B)** Stacked horizontal bar plot of significantly downregulated mitochondrial genes in NLFTau^m/h^ vs NLF. Genes were grouped into pathways extracted from the mouse MitoCarta 3.0 database. Bar length indicates gene count per pathway, with stacked segments representing sub-mitochondrial localisation. **(C)** Top mitochondrial DEGs from the NLFTau^m/h^ vs NLF comparison (Two-way ANOVA). Each point represents a biological replicate. *Post hoc* Tukey’s multiple comparisons test shown as ***p<0.001, **p< 0.01; main effect of Tau^m/h^ shown as †††† p<0.0001, †† p<0.01. **(D)** Volcano plot displaying DEGs in NLFTau^m/h^ vs NLF, derived from the SynGO 1.3 synaptic gene set. The dashed black line indicates an adjusted p-value threshold of 0.05, above which genes are considered significant. **(E)** Synaptic DEGs from the NLFTau^m/h^ vs NLF comparison (Two-way ANOVA). Each point represents a biological replicate. *Post hoc* Tukey’s multiple comparisons test shown as ***p<0.001, **p< 0.01,*p<0.05; main effect of Tau^m/h^ shown as †††† p<0.0001. **(F)** Schematic of significantly downregulated synaptic genes from the NLFTau^m/h^ vs NLF comparison. Gene annotations were derived from the SynGO 1.3 gene database. Created in BioRender (https://BioRender.com/dzj45zr). WT = 10, Tau^m/h^ = 8, NLF = 9, NLFTau^m/h^ = 3 mice.

To assess whether similar transcriptional alterations extended to synaptic pathways, a curated set of 1,674 synaptic genes were obtained from the SynGO 1.3 database^41^ and analysed using the same pipeline. In contrast to the mitochondrial dataset, differential expression within the synaptic gene set was more restricted, with only 8 DEGs identified in the NLFTau^m/h^ versus NLF comparison (Fig. 7D). Despite the smaller number of affected genes, these transcripts mapped to both pre- and post-synaptic compartments. For example, *Lrrk2* and *Gripap1* are implicated in vesicle dynamics and receptor trafficking, respectively, highlighting targeted perturbations in synaptic signalling pathways. Complete comparisons can be found in Supplementary Table S7. These transcriptomic findings are supported by the proteomic dataset, which showed enrichment in GO terms related to vesicle-mediated transport, synaptic exocytosis, and receptor-mediated endocytosis (Fig. 6C, 7E). Consistent with the mitochondrial analysis, transcriptional changes in synaptic genes were primarily associated with presence of human tau and further exacerbated by *App* mutations in NLFTau^m/h^ mice (Fig. 7F). Collectively, these findings highlight coordinated synaptic and mitochondrial dysregulation at both transcriptomic and proteomic levels, consistent with observations from other AD mouse models and human post-mortem studies^42–44^.

### Human-mouse comparisons identify shared AD pathways in NLFTau^m/h^ mice

We next investigated the translational relevance of mouse proteomic signatures by comparing them to those from published human post-mortem AD datasets^45^. To mimic the comparison of patients with AD to healthy controls, we compared NLFTau^m/h^ to WT mice, capturing molecular alterations arising from the combined effects of *App* mutations and human tau expression (Fig. 8A; complete list in Supplementary Table S2).

**Figure 8.**
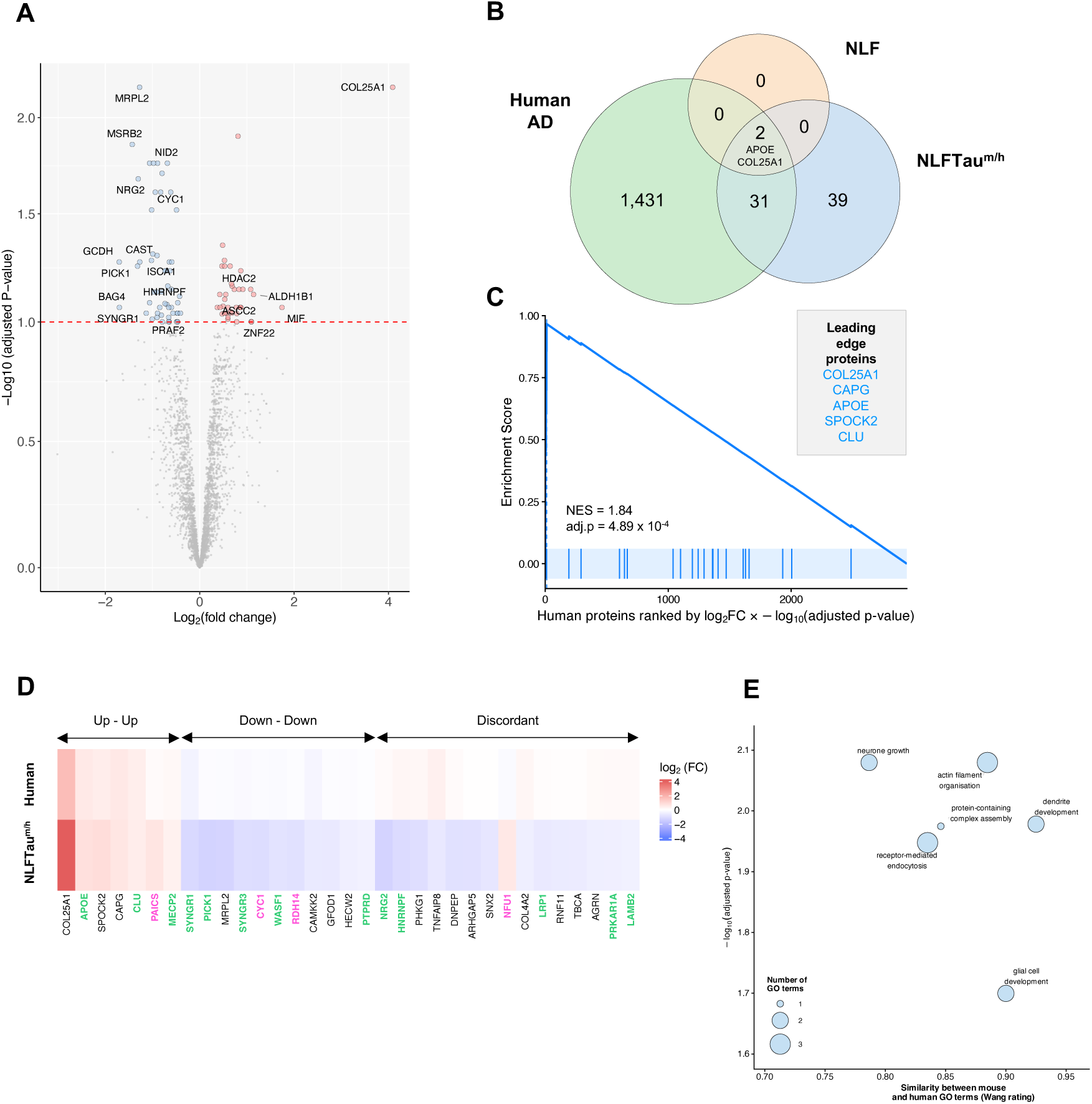
Conserved proteomic changes between NLFTau^m/h^ and human Alzheimer’s disease. **(A)** Volcano plots displaying DEPs in NLFTau^m/h^ vs WT. The dashed red line indicates an adjusted p-value of 0.1, above which proteins are considered significant. Labelled proteins are those with absolute log_2_(fold-change) > 1 **(B)** Venn diagram showing an overlap of shared DEPs between human AD, NLF and NLFTau^m/h^ mice. **(C)** Protein set enrichment analysis of upregulated NLFTau^m/h^ DEPs plotted against a human AD proteomic dataset ranked by log_2_(fold-change) weighted by statistical significance (–log_10_ adjusted p-value). Vertical tick marks along the x-axis show the location of individual NLFTau^m/h^ DEPs within the ranked protein list. **(D)** Heatmap showing log_2_(fold-change) of shared DEPs between human AD and NLFTau^m/h^, identified in **(B)**. Synaptic proteins are labelled below in green and mitochondrial proteins in pink. **(E)** Bubble plot illustrating semantic similarity between clusters of mouse BP GO terms and human GO terms in AD. Mouse GO terms were grouped into semantically related clusters, and each cluster was compared to human GO terms using the Wang similarity metric (mgoSim).

This analysis was concentrated on the NLFTau^m/h^ model because, at 24 months, NLF mice, consistent with their limited pathology, exhibited minimal differential expression compared to WT, thereby restricting meaningful cross-species comparisons (Fig. 5Aiv). In contrast, NLFTau^m/h^ mice displayed a substantially larger set of DEPs which overlapped with human AD datasets, allowing for more robust comparative analyses (Fig. 8B). When human proteins are ranked in order of fold change weighted by statistical significance from maximum increase through to maximum decrease, protein set enrichment analysis showed that upregulated DEPs in NLFTau^m/h^ mice were significantly enriched and concentrated toward the top of the human ranking (Fig. 8C). At the level of individual proteins, 33 DEPs were shared between NLFTau^m/h^ mice and human AD, of which 55% showed directional concordance (Fig. 8B, D). Notably, some proteins encoded by genes implicated in AD genome-wide association studies, including APOE, AGRN, and CLU, were differentially expressed in NLFTau^m/h^ mice^46^. In addition, a subset of overlapping proteins was associated with mitochondrial and synaptic pathways, including CYC1 and SYNGR1, consistent with the pathway-level disruptions identified in earlier analyses (Fig. 6B-C, 8D). Considering inevitable species differences but also, importantly, the different stages of AD represented in post-mortem human brains versus NLFTau^m/h^ mice at 24 months, differences in protein expression would be expected. Despite these unavoidable mismatches, there is clearly a notable overlap.

To investigate this further, we applied GO semantic similarity analysis to identify conserved biological processes across disease progression between mice and humans. Mouse GO terms were first grouped together into clusters, based on semantic similarity. Each cluster was then compared to human GO terms (Fig. 8E). This approach revealed convergent enrichment in processes such as neurone growth, receptor-mediated endocytosis, actin filament organisation, and glial cell development (full list of clustered mouse GO terms in Supplementary Table S8).

## DISCUSSION

Many studies have sought to clarify the relationship between Aβ accumulation and NFT formation. The most widely accepted hypothesis, proposed more than 30 years ago, suggests that Aβ aggregation initiates tau pathology and neuronal loss^47^. Despite the ‘amyloid cascade hypothesis’ being consistent with most of the existing data, it has not been demonstrated in any mouse model. This limitation has hindered efforts to define mechanistically how Aβ triggers or accelerates tau pathology, and how tau pathology ultimately leads to neurodegeneration and cognitive decline. Mouse models with Aβ pathology expressed alongside mutation-dependent tau pathology have been informative^5,6,48^. However, this approach implicitly assumes that downstream cellular mechanisms, and, by extension, therapeutic targets and diagnostic biomarkers, are conserved, regardless of whether Aβ and tau pathologies arise sequentially or in parallel through distinct routes of onset. To date, there is currently little empirical evidence to distinguish between these possibilities. This incomplete understanding of disease progression has limited our avenues for therapeutic development and diagnostic innovation. Most disease-modifying strategies have adopted an Aβ-centric approach, aiming to reduce Aβ production or promote plaque clearance. Monoclonal antibodies targeting Aβ, such as lecanemab and donanemab, have demonstrated the capacity to reduce amyloid burden and modestly slow cognitive decline, representing a significant milestone in the field^49,50^. This implies that the Aβ burden is clearly contributing to AD-related cognitive decline but whether it is directly causal is open to question.

Here, we present a series of mouse models that overcomes a major limitation of previously available models. The most novel of these models, the NLFTau^m/h^ cross, exhibits a delayed initiation of NFT formation arising from the interaction between ageing, Aβ accumulation and human tau expression. Most existing models rely on rapid plaque accumulation at early ages, compressing disease timelines for reasons of cost and convenience^19,28,51,52^. In contrast, the NLF background produces a slower trajectory of Aβ deposition, aligning more closely with the preclinical stages of AD^19,20^. Given that age is the strongest risk factor for AD^53^, this extended timeframe allows interrogation of ageing-dependent disease mechanisms that are otherwise missing in accelerated models. Moreover, the use of knock-in technology avoids artefacts associated with transgenic overexpression.

A defining key feature of adult humans, relevant to AD versus other tauopathies, is the presence of both 3R and 4R tau isoforms at an approximate ratio of 40:60^22,54^; a feature recapitulated in this model. Fully humanised *MAPT* mouse models often exhibit a relative enrichment of 3R over 4R tau, a shift that may limit the formation of NFT pathology^11,55^.

An early step in NFT formation is tau hyperphosphorylation, which destabilises tau-microtubule interactions, promoting somatodendritic mislocalisation and/or aggregation^56,57^. Tau hyperphosphorylation as seen in human AD ^23,26^ is mimicked in NLFTau^m/h^ mice at 24 months. Interestingly, with progression to extreme old age, both NLF and Tau^m/h^ mice also develop similar tau pathology. Age-related increases in hippocampal p-tau have been linked to synaptic and mitochondrial dysfunction^58^, key hallmarks of p-tau-associated neurodegeneration^57,59,60^. Consistent with this, genotypes expressing human tau show coordinated reductions in mitochondrial and synaptic gene and protein expression, which are further exacerbated by Aβ. These findings suggest that human tau expression introduces an early, network-level vulnerability, likely reflecting fundamental differences between human and mouse tau, that predisposes neurones to later tau pathology. This is likely further exacerbated by our observation of age-dependent increases in chromatin condensation across genotypes. Overall, this stresses the essential element of advanced old age in these models.

Thus, while either Aβ or human tau alone is sufficient to initiate NFTs in extreme old age, their interaction accelerates this process, leading to an earlier onset when both are present. The appearance of somatic p-tau at 28 months in Tau^m/h^ mice, in the absence of Aβ, mirrors pathological features of primary age-related tauopathy, where p-tau pathology arises independently of Aβ in the ageing brain^61,62^.

The question arises as to how Aβ pathology accelerates NFT formation. The progressive accumulation of p-tau within cored plaques and its correlation with NFT formation in individual mice suggests a causal link between initial Aβ-associated p-tau, putatively in axonal terminals, and the eventual development of NFTs, as previously proposed^63^. This interpretation is further supported by human post-mortem studies, which show that clinical dementia is associated with a higher prevalence of cored, neuritic plaques, whereas cognitively normal amyloid positive individuals predominantly exhibit diffuse plaques^64,65^. Moreover, we have previously shown that plaque structural maturation correlates with increased neurotoxicity and synaptic damage^66^. The greater susceptibility of human tau to misfolding and aggregation, relative to murine tau, may enhance its vulnerability to hyperphosphorylation within cored plaques, thereby promoting earlier somatodendritic p-tau mislocalisation, as observed in 24-month-old NLFTau^m/h^ mice^11,67,68^.

At 24 months, the NLFTau^m/h^ model demonstrates greater proteomic concordance with human AD than the NLF model, highlighting the importance of human tau in shaping a disease-relevant molecular landscape. By recapitulating Aβ- and age-dependent tau hyperphosphorylation, NLFTau^m/h^ satisfies key criteria for an improved AD mouse model, opening future possibilities for deciphering the molecular mechanisms linking these interacting factors. Although this still represents an early phase of AD before full tau pathology and neurodegeneration, availability of a model that at least initiates NFT formation without mutations in *MAPT* will allow testing of drugs that could break the link between Aβ and tau and study the progression of tau pathology. Combining this model with genetic or environmental risk factors may accelerate disease progression and permit investigation of pathology across the full spectrum. The NLFTau^m/h^ mouse can now be used to guide the development of new treatments aimed at preventing or slowing clinical manifestations of the disease, with every chance that findings in mice will translate to successful treatments in humans.

## METHODS

### Animals

All experiments were carried out in accordance with the UK Animal (Scientific Procedures) Act, 1986, following local ethical review. This study compared mixed males and female, age-matched mice of 4 genotypes:

1. *App^NL-F/NL-F^* knock-in mice (NLF);
2. *App^NL-F/NL-F^* mice harbouring heterozygous humanised knock-in *MAPT* (NLFTau^m/h^);
3. heterozygous humanised knock-in *MAPT* (Tau^m/h^) and
4. C57BL/6J wild type (WT).

*Mapt* knock-out mice were used as negative controls and were kindly provided by Prof. Karen E. Duff. Same sex littermates were group-housed (2-5 mice) at the UCL Biological Services Unit under a 12-hour light/dark cycle with *ad libitum* food and water supply, at a controlled temperature and humidity. The NLF and *MAPT* knock-in mice described are available from RIKEN BRC (https://web.brc.riken.jp/en).

### Genotyping

Genotyping was performed by Transnetyx genotyping services (Cordova, TN, USA), using ear or tail tissue.

### Tissue Extraction

Animals were decapitated and brains were rapidly extracted on ice and hemisected. One hemisphere was drop-fixed in 4% paraformaldehyde at 4°C overnight and then stored in 30% sucrose, 0.02% sodium azide phosphate-buffered saline (PBS; 137 mM NaCl, 2.7 mM KCl, 10 mM Na_2_HPO_4_, 1.8 mM KH_2_PO_4_). The other hemisphere was dissected to extract hippocampus, cortex and cerebellum, which were snap-frozen on dry ice at –80°C for RNA and protein extraction.

### Western blot

Cortical mouse brain tissue was homogenised in lysis buffer (10 mM Tris-HCl (pH 7.4), 0.8 M NaCl, 1 mM EDTA, 10% sucrose, cOmplete mini protease inhibitor cocktail (Roche Diagnostics #11836153001), centrifuged at 12,000 *g* for 5 minutes at 4°C. For detection of tau isoforms, brain extracts were dephosphorylated with 0.8x MnCl_2_ and protein metallophosphatases, and 16,000 units/ml lambda protein phosphatase at 30°C for 3 hours (New England Biolabs #P753S).

Samples were run on a 10% Bis-Tris 15-well mini 1.5 mm NuPAGE system at 160 V for 4 hours, followed by an overnight PVDF transfer at 15 V. Membranes were blocked for 1 hour at room temperature (RT) in 5% non-fat milk, 0.1% Tween 20 tris buffered saline (TBST; 20 mM Tris-HCl, 137 mM NaCl, pH = 7.6) followed by an overnight incubation at 4°C with 1:1500 anti-mouse Tau-5 antibody (Invitrogen #AHB0042) in superblock blocking buffer. The membranes were washed three times in TBST, incubated with 1:2000 horseradish peroxidase goat anti-mouse secondary antibody (Agilent #P044701-2) for 1 hour at RT in TBST with 5% (w/v) non-fat milk, washed six times in TBST and imaged using the BioRad Chemidoc imaging system. Ratio of 4R:3R peptides was quantified using Image Lab.

### Mass spectrometry analysis targeting tau peptides

#### Brain extraction, immunoprecipitation and protein digestion

Cortical brain tissue was homogenised in TBS (containing cOmplete mini protease inhibitor cocktail) for 2 minutes at 200 Hz using Tissue Lyser II (Qiagen), centrifuged at 31,000 *g* for 1 hour at 4°C. Protein concentration was measured on the supernatant using the DC Protein Assay kit (Bio-Rad). The analysis of tau protein, entailing the steps of immunoprecipitation, proteolytic digestion and mass spectrometry analysis, was carried out as described previously^22^. Briefly, 20 µg of total protein from brain extracts was immunoprecipitated using 50 µl M-280 Dynabeads (Invitrogen) coated with 4 µg 77G7 anti-tau antibody (MTBR epitope 316-355, BioLegend) per sample. The antibody was chosen to bind both human and mouse tau protein. To each sample, 10 µl (containing 133 fmol each) of tau 0N3R, 1N4R, and 2N4R isoforms standards, uniformly labelled [U-^15^N], were also added. Following overnight incubation at 4°C, samples were washed, and bound proteins were subsequently eluted with 0.5% formic acid. Eluted samples were then dried in a vacuum centrifuge and stored at -80°C. Proteolytic digestion was carried out overnight at 37°C by reconstituting the samples in 50 mM ammonium bicarbonate containing 100 ng trypsin (V5111, Promega) and 100 fmol of labelled phosphorylated peptides specifically labelled at lysine [^13^C,^15^N-Lys] or arginine [^13^C,^15^N-Arg] residues, see Table S1. The next day, trypsination was stopped with the addition of 2 µl 10% formic acid. Samples were dried in a vacuum centrifuge and stored at -80°C pending mass spectrometry analysis.

#### Liquid chromatography-mass spectrometry (LC-MS)

Immunoprecipitated and trypsinated samples were reconstituted into 7 µl of 8% formic acid/8% acetonitrile in water of which 6 µl were injected. LC-MS analysis was carried out using a Dionex 3000 system (Thermo Fisher Scientific) coupled to an electrospray ionisation hybrid quadrupole-orbitrap mass spectrometer (Q Exactive, Thermo Fisher Scientific). LC separation was performed at a flow of 300 nl/minute using a gradient. The mass spectrometer was set to acquire in the m/z range 350–1400 in data-dependent mode using higher-energy collision-induced dissociation for ion fragmentation. Details of analysis have been previously described ^22^.

#### Processing of LC-MS data

Database searches of acquired peaks used PEAKS Studio Xpro (Bioinformatics Solutions) and Mascot Daemon v2.6.1/Mascot Distiller v2.6.3/ Mascot database search software v2.6.1 (Matrix Science). Searches were made against both UniProt and a custom-made tau-only database containing both human and mouse sequences. Quantitative analysis was performed using Skyline software v20.1.0.31 (MacCoss Lab) using the first two isotopes of the precursor ions. Search data was used to confirm the identities of peaks detected by Skyline and every peak was manually inspected. Data are shown as light-to-heavy peptide ratios. Table S1 shows the peptides included for final quantification.

### Immunohistochemistry

Thirty μm free-floating formaldehyde-fixed sections were cut transverse to the long axis of the hippocampus using a frozen sledge microtome (Leica SM2010R) and collected into PBS with 0.02% sodium azide. Sections were washed in PBS for 10 minutes, followed by antigen retrieval in 10 mM sodium citrate buffer (pH 9.0) at 80°C for 30 minutes. Tissues were permeabilised in three 10-minute 0.3% Triton-X 100 PBS (PBST) washes and then incubated in blocking buffer (3% normal goat, 0.3% PBST) for 1 hour at RT. The sections were subsequently incubated overnight at 4°C with appropriate primary antibodies in blocking buffer; 1:1000 rabbit anti-phosphotau-serine396, (pS396, Invitrogen #44-752G) and 1:500 mouse anti-amyloid, 1-16 (6E10, Biolegend #803004).

The next day, sections were washed three times in 0.3% PBST for 10 minutes each, followed by a 2-hour incubation with appropriate secondary antibodies in blocking buffer in the dark at RT; 1:1000 goat anti-rabbit Alexa Fluor 647 (Invitrogen #A21244) and 1:1000 goat anti-mouse Alexa Fluor 594 (Invitrogen #A11032). Subject to three 10-minute 0.3% PBST washes, sections were incubated with a combination of two luminescent conjugated oligothiophenes (LCOs, 1:500 tetrameric and 1:1500 heptameric formic thiophene acetic acid; q-FTAA and h-FTAA, respectively) in the dark at RT for 30 minutes. Finally, nuclei were counterstained with 1:10000 4′,6-diamidino-2- phenylindole (DAPI) for 5 minutes at RT.

### Fluorescent Imaging

All imaging was performed on the Zeiss LSM 880 confocal microscope with a 32-channel GaAsP spectral detector. For LCO analysis, q- and h-FTAA dyes were excited by a 458 nm argon laser. Using reference emission spectra of the two dyes obtained by hyperspectral imaging in lambda mode, linear unmixing was subsequently performed to obtain separate images for q- and h-FTAA in the multi-stained samples, as described previously^66^.

Confocal photomicrographs of hippocampal tissue, multi-stained with q-FTAA and h-FTAA, were acquired as z-stack images (3-μm intervals) using a Plan-Apochromat 20× objective (NA 0.8). Additionally, the same samples were stained with DAPI, pS396, and 6E10, and imaged as widefield photomicrographs using the same objective. These images were then merged into a single file for comprehensive analysis. Images of DAPI in the CA1 and CA3 pyramidal layer were obtained using a Plan-Apochromat 40× oil objective (0.8 NA) under constant laser power, master and digital gain, and offset settings.

### Immunohistochemical analysis

**I**mmunohistochemical staining analyses were performed using custom-made semi-automated macros on ImageJ (as described below):

#### Plaque load analysis

The hippocampal region for each image was manually selected and thresholded to adjust for individual background differences, as previously described^69^. Particles less than 10 μm^2^ were excluded from analysis. Plaques were subsequently classified as either q-FTAA+ve, h-FTAA+ve, Aβ+ve (cored plaques) or q-FTAA-ve, h-FTAA+ve, Aβ+ve (fibrillar plaques). (q-FTAA-ve, h-FTAA-ve, Aβ+ve (diffuse Aβ) were excluded from this analysis.)

#### Protein analysis around plaques

For each cored and fibrillar plaque identified within the selected hippocampal region of interest (ROI), concentric contours, preserving the original shape of each plaque, were drawn outwards with increasing radial increments (radii from plaque edge (μm); 0, 5, 10, 20, 30). Where concentric contours of cored and fibrillar plaques overlapped, the ROI in this overlapping region was omitted from further analysis. It is noteworthy that while all LCO plaques are also positive for Aβ, not all Aβ staining is positive for LCO. To exclusively analyse the spatial effect of Aβ on protein intensity, concentric contours, as described above, were drawn for Aβ+ LCO- regions. ROIs with overlapping contours between Aβ+ and LCO+ plaques were excluded from further analysis. Subsequently, mean protein fluorescence intensity across the plaque and within each concentric contour was calculated as a fold-change compared to the 20-30 μm ring around the respective plaque type.

#### Protein analysis within hippocampal pyramidal cells

In the hippocampal CA1 and CA3 pyramidal cell layer, concentric contours were drawn around each identified DAPI nucleus to maintain its original shape, extending 3.5 μm away from the DAPI edge to delineate cytoplasmic regions as ROIs. Any overlapping regions were merged as a single larger ROI. Protein fluorescence intensity within these ROIs was calculated as fold-change from mean protein fluorescence intensity in the axonal region. Analysis was verified independently by two members of the lab, each blind to genotype.

#### DAPI-positive cell counts

To quantify cell densities, three non-overlapping confocal photomicrographs were acquired for each technical replicate from the hippocampal CA1 and CA3 pyramidal cell layers. Within each image, a rectangular ROI was defined, and DAPI-positive nuclei within the ROI were counted. Objects touching the boundaries of the rectangular ROI were included only on two adjacent sides.

#### Chromatin analysis within neuronal nuclei

DAPI images were processed in Ilastik^70^ (version 1.4.1) using the pixel- and object-classification workflows to segment chromatin based on machine-learning classification. The resulting masks were exported as binary images. To quantify chromatin coverage, ROIs corresponding to DAPI+ve nuclei were generated, and the area occupied by chromatin within each ROI was measured.

### Kaplan-Meier survival curve

The Kaplan-Meier curve was plotted in GraphPad Prism (version 10.6.1). Survival time was defined as the number of days from birth to event. Event status was coded as a binary variable (0 = alive), 1 = death. On any day, all live mice were included in the analysis. If a mouse died at a predefined endpoint or was culled for reasons unrelated to neurodegeneration, the event of its death was excluded from analysis. An event (status = 1) was assigned to mice that reached a neurodegenerative-like endpoint characterised by hunched posture, sustained weight loss, piloerection, and akinesia, or were found dead during the study. Cox proportional hazards regression models were used to assess the effects of APP and Tau^m/h^ genotype. Hazard ratios with corresponding 95% confidence intervals were estimated.

### Untargeted LC-MS based proteomics

#### Protein extraction and digestion

For the MS analysis, cortical brain sections were solubilised with RIPA buffer (50 mM Tris-HCl (pH 7.6), 150 mM NaCl, 1% NP-40, 1% sodium deoxycholate, 0.1% SDS, 1 x protease inhibitor cocktail (Thermo Fisher Scientific #78443) followed by ultrasonication in Bioruptor Pico Plus (10 cycles, medium frequency). Proteins were precipitated using chloroform/methanol protein precipitation followed by denaturation, reduction, alkylation, enzymatic digestion and peptide cleanup. Methanol and chloroform were added in a 4:1:3 ratio (sample: methanol: chloroform), followed by vortex and centrifugation at high speed to separate phases. The aqueous layer was discarded, and the protein pellet was washed with methanol and briefly air-dried. The pellet was then resuspended in 100 µl of 6 M guanidine hydrochloride (Sigma-Aldrich #G4505) and incubated at 95 °C for 10 minutes to ensure complete solubilisation. Disulfide bonds were reduced by adding 0.5 µl of 1 M DTT (DOT Scientific Inc. #DSD11000) followed by incubation at 56 °C for 20 minutes. After cooling to RT, 3 µl of 500 mM iodoacetamide (IAA, Sigma-Aldrich #I1149) was added to alkylate reduced cysteine residues. Samples were incubated in the dark at RT for 20 minutes. Excess IAA was quenched by adding 5 µl of 1 M DTT and incubating for 15 minutes at RT. Then, the samples were digested with mass spec grade trypsin/Lys-c mix (Promega #V5073) overnight at 37°C. The reaction was subsequently stopped by acidification with 1% formic acid and desalted using C18 spin columns (Thermo Scientific #89852). Eluates were dried using a vacuum concentrator and stored at −80 °C.

#### Tandem mass tag-MS sample preparation

A 16-plex tandem mass tag (TMT) was performed on the MS samples. The TMT preparation was performed as previously described^71^. The dried peptides were resuspended in 100 mM HEPES, pH 8.5. Micro BCA assay (Fisher Scientific #PI23235) was subsequently performed to determine the concentration of peptides. Eighty μg of peptide from each sample was then used for isobaric TMT labelling, following manufacturer’s instructions (0.4 mg, dissolved in 40 μL anhydrous acetonitrile; ThermoFisher Scientific #A44520). After incubating for 2h at RT, the reaction was quenched with 5% (v/v) hydroxylamine to 0.3%. Isobarically labelled samples were then combined in equal amounts and quickly desalted with C18 HyperSep column (Thermo scientific #60108-302). The combined TMT labelled peptide samples were fractionated into 8 fractions: 5.0%, 10.0%, 12.5%, 15.0%, 17.5%, 20.0%, 22.5%, 25.0% and 50% of ACN in 0.1% triethylamine solution, using high pH reversed-Phase columns (ThermoFisher Scientific #84868). Peptide solutions were dried, stored at −80 °C, and reconstituted in LC-MS Buffer A (5% acetonitrile, 0.125% formic acid) for LC-MS/MS analysis.

#### LC-MS/MS

Dried peptide samples were resuspended in 20 µl of Buffer A (94.875% H₂O, 5% acetonitrile (ACN), and 0.125% formic acid). A total of 3 µg of peptide was injected via autosampler and loaded onto a PepMap 100 C18 nanoViper trap column (75 µm × 2 cm, Thermo Fisher Scientific) using an UltiMate 3000 HPLC system. Peptides were then separated on an analytical PepMap RSLC C18 column (Thermo Fisher Scientific) coupled to a stainless-steel emitter tip and analysed using a Nanospray Flex ion source operating at 2,000 V. The chromatographic run was 4.5 hours in total with the following profile of Buffer B: 2% for 7 mins, 2 – 7% for 1 min, 7 – 10% for 5 mins, 10 – 25% for 160 min, 25 – 33% for 40 min, 33 – 50% for 7 min, 50 – 95% for 5 min, 95% for 15 min, then back to 2% for the remaining 30 min. Peptides were analysed on an Orbitrap Fusion mass spectrometer (Thermo Fisher Scientific) using data-dependent acquisition. We used a multiNotch MS3-based TMT method to analyse all the TMT samples^72,73^. MS1 survey scans were acquired in the Orbitrap at a resolution of 60,000, over a scan range of 300–1,500 m/z. MS parameters were: ion transfer tube temperature, 300 °C; Easy-IC internal mass calibration; default charge state, 2; automatic gain control (AGC) target, 2×10⁵; maximum injection time, 50 ms; microscans, 1; and S-lens RF level, 60. Data were collected in positive ion mode with centroid format. MIPS was enabled. Charge states from 2–6 was selected for fragmentation; unassigned precursors were excluded. Dynamic exclusion was enabled with a repeat count of 1, exclusion duration of 30 s (high) and 45 s (low), and a ±10 ppm mass tolerance.

For MS/MS (MS2) acquisition, the top 20 most intense precursor ions were selected for fragmentation. Precursor isolation was performed with a 1.6 m/z window using the quadrupole. Higher-energy collisional dissociation was employed with a normalised collision energy of 30%. Fragment ions were detected in the ion trap using rapid scan rate, with a maximum injection time of 75 ms and an AGC target of 10,000. Q value was set to 0.25, and ion accumulation used all available parallelisable time.

#### Processing of LC-MS/MS data

Protein identification/quantification and analysis were performed with Integrated Proteomics Pipeline - IP2 (Bruker, Madison, WI. http://www.integratedproteomics.com) using ProLuCID Ver 6.5.5, DTASelect2, Census and Quantitative Analysis. Spectrum raw files were extracted into MS1, MS2 and MS3 files using RawConverter (http://fields.scripps.edu/downloads.php). The tandem mass spectra were searched against combined_HumanTau_Mouse_withHumanSwedishAPP_Human_mouse_02-07-2024.fasta, matched to sequences using the ProLuCID/SEQUEST algorithm with 20 ppm peptide mass tolerance for precursor ions and 600 ppm for fragment ions. ProLuCID searches included all fully and half-tryptic peptide candidates that fell within the mass tolerance window and had with no miscleavages. The search parameters included static modifications for cysteine carbamidomethylation (+57.0215 Da) and TMT pro-labelling of lysine and peptide N-termini (+304.2071 Da). Differential modifications were allowed for the phosphorylation of serine, threonine, and tyrosine (+79.9663 Da), with a maximum of two internal variable modifications per peptide. Peptide/spectrum matches (PSMs) were assessed in DTASelect2 using the cross-correlation score (XCorr), and normalised difference in cross-correlation scores (DeltaCN). Peptide probabilities and false-discovery rates (FDR) were calculated based on a target/decoy database containing the reversed sequences of all the proteins appended to the target. For each biological replicate, proteins identified had an FDR of ≤1% at the protein level. Each protein identified was required to have a minimum of one peptide of minimal length of six amino acid residues.

#### Data normalisation

To normalise the data, individual protein abundances were first log_2_-transformed, then each value was centred by subtracting the sample median and finally shifted by adding the global median. Proteins with more than 50% of missing values within any one group were excluded from subsequent analysis.

### Proteomic data analysis

#### Differential expression analysis

A linear model was fitted using the limma package for proteome-wide differential abundance analyses, followed by empirical Bayes moderation to improve variance estimation^74^. Differentially expressed genes proteins (DEPs) were defined using an adjusted p-value < 0.1, reflecting the reduced statistical power associated with limited sample size and milder pathology, which may attenuate detectable changes in bulk proteomics.

#### Gene ontology

Gene ontology (GO) analysis was performed using the clusterProfiler package^75^. Overrepresentation analysis was conducted to identify GO terms within the biological process, molecular function, and cellular component categories that were significantly enriched among the selected proteins within the background proteome using the org.Mm.eg.db database. Multiple testing correction was applied using the Benjamini–Hochberg method, with significance defined as p < 0.05. Semantic similarity analysis of GO terms across mouse and human datasets was performed using Wang’s method implemented in the GOSemSim package to quantify functional relatedness^76^.

#### Weighted protein co- expression network analysis

The weighted protein co-expression networks were constructed using the WGCNA package^77^. Modules were identified using dynamic tree applied to the topological overlap matrix with the following parameters: soft threshold power = 6 (R² ≥ 0.85 for approximate scale-free topology), corType = bicor, networkType = signed, TOMType = signed, TOMDenom = mean, deepSplit = 2, minModulesize = 30, mergeCutHeight =0.25, pamRespectsDendro = TRUE, reassignThreshold = 0.05, and minKMEtoStay = 0.30. Ten modules were identified in the proteomics dataset, along with a grey module representing unassigned proteins. Eigenvalues, the first principal component of each module, were then calculated. Pearson correlations were used to determine module eigenvalue-based connectivity (kME) for each protein.

#### Protein-protein interaction network

Intramodular correlation networks were constructed using kME values and exported to Cytoscape (version 3.10.3) for visualisation. Only top 30 nodes per module with kME value > 0.70 were displayed.

#### Protein set enrichment analysis

Mouse proteins were mapped to their corresponding human orthologs using Ensembl BioMart. Pre-ranked protein set enrichment analysis was then performed in R using the fgsea package. Human proteins were ranked using a composite metric defined by log_2_(fold change) x - log_10_(adjusted p-value), integrating both effect size and statistical significance. Mouse-derived proteins were then tested for enrichment against this human AD proteomic dataset, based on previously published data^45^. Enrichment score (ES) was calculated using the GSEA running-sum statistic, with statistical significance assessed by permutation testing.

### RNA Extraction and Quantification

Frozen hippocampal tissue was homogenised in QIAzol RNA lysis reagent using a Polytron PT 3000 homogeniser at 7000 rpm for 30 seconds. Chloroform was added to facilitate phase separation, followed by centrifugation at 12,000 x *g* for 15 minutes. RNA was isolated and subjected to DNA digestion using the miRNeasy Mini Kit according to the manufacturer’s instructions. RNA purity and concentration were determined spectrophotometrically by measuring A260/A280 ratios. RNA samples were stored at -80°C till further use. RNA was processed by Eurofins Genomics (Eurofins Genomics Europe Sequencing GmbH, Germany) for RNA sequencing using the NextSeq 2000 RNA sequencer.

### Transcriptomic Analysis

#### Data processing and normalisation

Raw counts matrices were imported into R (version 4.5.2) for downstream analysis. Lowly expressed genes were filtered using the *filterByExpr* function in edgeR. Gene annotations were retrieved from the Ensembl BioMart database, and analyses were restricted to protein-coding genes. Library size normalisation was performed using the trimmed mean of M-values (TMM) method, followed by normalisation to housekeeping genes (*Actb* and *Actg1*) to control for technical variation. Mitochondrial and synaptic genes were extracted from the MitoCarta 3.0 and SynGo 1.3 dataset respectively^40,41^.

#### Differential expression analysis

Differential expression analysis for mitochondrial and synaptic genes was conducted using the limma-voom pipeline, followed by empirical Bayes moderation to improve variance estimation^74^. To control for multiple testing, p-values were adjusted using the Benjamin-Hochberg correction. Genes were considered significantly differentially expressed at adjusted p-value of < 0.05.

### Statistical Analysis

Analysis was performed using R Statistical Software (version 4.5.2; R Core Team 2021), GraphPad Prism (version 10.6.1) or SPSS (version 31.0.0.0). Samples sizes represent number of animals and any technical replicates obtained from a single animal were averaged for that animal. Analysis of variance or generalised linear mixed model was applied as appropriate. Investigators were blinded to genotype throughout the experiments, unless blinding was not feasible.

## Data and materials availability

Datasets generated and/or analysed for this study will be made available from the source data or made publicly available upon publication.

## Code availability

All coding performed is publicly available in: https://github.com/snehadesai1029/Desai-et-al.-2026

## ACKNOWLEDGEMENTS

This work was primarily supported by the Cure Alzheimer’s Fund (FAE, J.Hardy, J.Hanrieder, HZ, GB, EC). SD was supported by the Research Excellence Scholarship from University College London. Other contributions were from: The National Institutes of Health (R01 AG078796 J.Hanrieder, FAE, JNS and S10OD032464 JNS); the Swedish Research Council (#2023-02796, J.Hanrieder); Alzheimerfonden (AF-1032753, J.Hanrieder); the Swedish Brain Foundation (Hjärnfonden, FO2025-0126, J.Hanrieder; #FO2022-0270, HZ); Swedish State Support for Clinical Research (#ALFGBG- 1006922, J.Hanrieder; #ALFGBG-71320,HZ), Åhlén-stiftelsen (JH, GB); Stiftelsen för Gamla Tjänarinnor (#2025-329, J.Hanrieder, HZ, GB); Stohne’s Stiftelse (JH, HZ, GB). J.Hardy is supported by the Dolby Foundation. HZ is a Wallenberg Scholar and a distinguished Professor at the Swedish Research Council supported by grants from the Swedish Research Council (#2023-00356, #2022-01018 and #2019-02397); the European Union’s Horizon Europe research and innovation programme under grant agreement No 101053962, the Alzheimer Drug Discovery Foundation (ADDF), USA (#201809-2016862), the AD Strategic Fund and the Alzheimer’s Association (#ADSF-21-831376-C, #ADSF-21-831381-C, #ADSF-21-831377-C, and #ADSF-24-1284328-C), the European Partnership on Metrology, co-financed from the European Union’s Horizon Europe Research and Innovation Programme and by the Participating States (NEuroBioStand, #22HLT07), the Bluefield Project, the Olav Thon Foundation, the Erling-Persson Family Foundation, Familjen Rönströms Stiftelse, Familjen Beiglers Stiftelse, the European Union’s Horizon 2020 research and innovation programme under the Marie Skłodowska-Curie grant agreement No 860197 (MIRIADE), the European Union Joint Programme – Neurodegenerative Disease Research (JPND2021-00694), the National Institute for Health and Care Research University College London Hospitals Biomedical Research Centre, the UK Dementia Research Institute at UCL funded by MRC (UKDRI-1003), and an anonymous donor. Åhlén-stiftelsen, Stiftelsen för Gamla Tjänarinnor and Stohnes Stiftelse. TS is suppored by JST Moonshot R&D Program (JPMJMS2024) and AMED (Grant Numbers JP24wm0625303).

We would like to thank Prof. Peter Nilsson and Prof. Per Hammarström for providing us with the q-FTAA and h-FTAA fluorophores, that were developed in their lab at Linköping University. We also thank RIKEN BRC for providing us the *App*^NL-F/NL-F^ and humanised *MAPT* knock-in mice. In addition, the authors thank David Kinnon for conducting independent analyses that supported the methods developed in this manuscript.

## AUTHOR CONTRIBUTIONS

Conceptualisation: SD, FAE, DMC

Methodology: SD, SB, AA, JIW, DAS, TS, TCS

Investigation: SD, JL, EC, GB, KG, TT, NS

Analysis: SD, JL, AB, GB, HBH

Visualisation: SD

Funding acquisition: FAE, J.Hanrieder, J.Hardy, JNS

Supervision: FAE, J.Hanrieder, DMC

Writing – original draft: SD, FAE

Writing – review & editing: All authors

