## Supplementary Figures for "Age- and amyloid-β-dependent initiation of neurofibrillary tau tangles: NLFTau^m/h^, an improved mouse model of Alzheimer’s disease without mutations in *MAPT*"

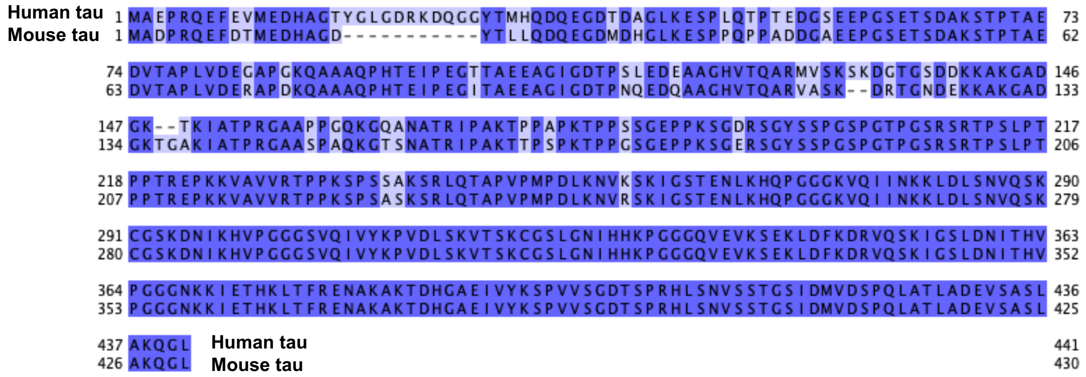


**Figure S1. Comparison of human and mouse 2N4R tau protein isoform sequence.** Amino acids highlighted in dark blue are identical between human and mouse tau, while light blue indicate differences. Dashed lines show regions where amino acids are missing in one of the sequences. Numbers along the sides serve as reference points for amino acid positions. Sequence alignment was made using Jalview.

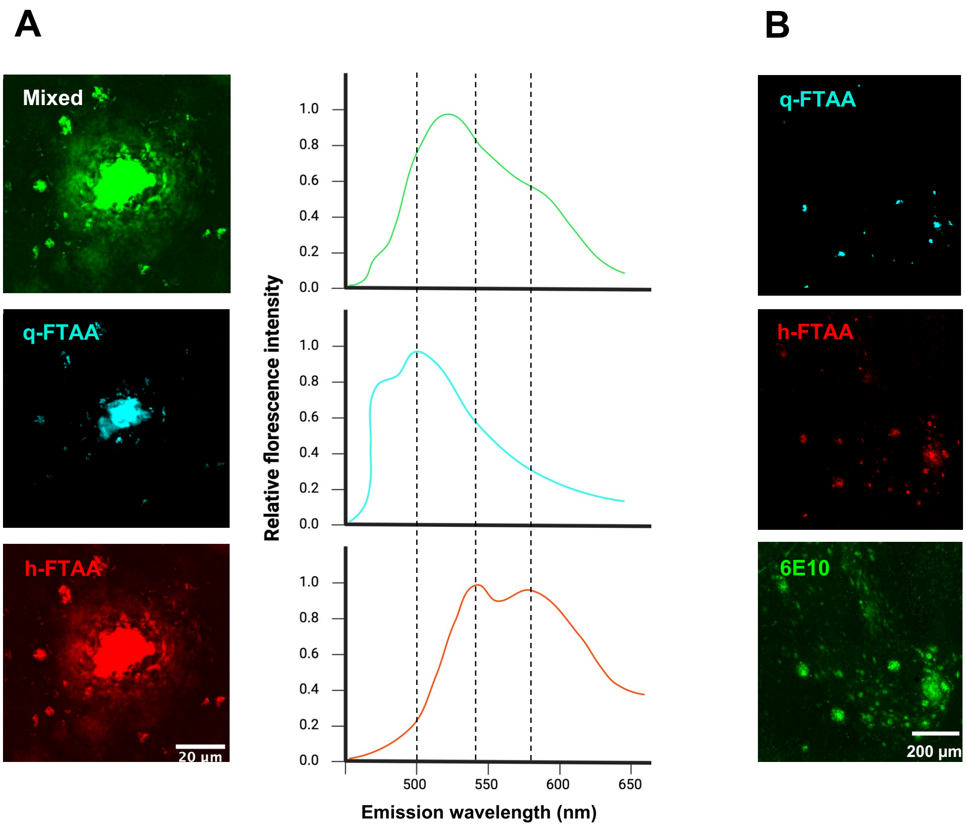


**Figure S2. Plaques can be classified according to their amyloid content using conformation-sensitive LCOs.**

**(A)** Each LCO exhibits a distinct spectral emission profile, identifiable with a hyperspectral detector in lambda mode (405 – 750 nm) for each pixel when stimulated at 458 nm. q-FTAA shows a blue-shifted peak at 500 nm, while h-FTAA displays red-shifted peaks between 540 and 580 nm. Linear unmixing allows for the separation of mixed LCO signals into distinct q-FTAA and h-FTAA channels. Parts of the panel created in Biorender (https://BioRender.com/y7a3uue) **(B)** An overview of q-FTAA, h-FTAA and 6E10 Aβ staining distribution in the hippocampus.
