## Supplementary Table 1 for "Age- and amyloid-β-dependent initiation of neurofibrillary tau tangles: NLFTau^m/h^, an improved mouse model of Alzheimer’s disease without mutations in *MAPT*"

| Measured peptide | Phosphorylation site | Peptide Sequence |
|---|---|---|
| 195-209 | NA | SGYSSPGSPGTPGSR |
| 212-221 | NA | TPSLPTPPTR |
| 243-254 | NA | LQTAPVPMPDLK |
| 275-286 (3R) | NA | VQIVYKPVDLSK |
| 282-290 (4R) | NA | LDLSNVQSK |
| 354-369 | NA | IGSLDNITHVPGGGNK |
| 396-406 | NA | SPVVSGDTSPR |
| 407-438 | NA | HLSNVSSTGSIDMVDSPQLATLADEVSASLAK |
| 175-190 p1 | T181 | TPPAPK**T**PPSSGEPPK |
| 195-209 p1 | S199 | SGYS**S**PGSPGTPGSR |
| | S202 | SGYSSPG**S**PGTPGSR |
| 212-221 p1 | T217 | TPSLP**T**PPTR |
| 225-234 p1 | T231 | KVAVVR**T**PPK |
| 386-406 p1 | S396 | TDHGAEIVYK**S**PVVSGDTSPR |
| 386-406 p2 | S396 + S404 | TDHGAEIVYK**S**PVVSGDT**S**PR |
| 396-406 p1 | T403 | SPVVSGD**T**SPR |
| | S404 | SPVVSGDT**S**PR |

**Table S1. Quantified tau peptides: phosphorylated and non-phosphorylated.**
Phosphorylation sites within specific peptide sequences are shown in bold. Phosphorylated peptide standards were labelled at lysine [$^{13}C_6$,$^{15}N_2$-Lys] or arginine [$^{13}C_6$,$^{15}N_2$-Arg] residues. All non-phosphorylated peptide standards were uniformly labelled with [$U$-$^{15}N$].
